# Tim-1 and Tim-4 mediate entry of the human Kaposi’s sarcoma-associated herpesvirus and the related rhesus monkey rhadinovirus

**DOI:** 10.1101/2023.10.10.561662

**Authors:** Stefano Scribano, Sarah Schlagowski, Shanchuan Liu, Thomas Fricke, Xiaoliang Yang, Frank Neipel, Anna K. Großkopf, Bojan F. Hörnich, Marija Backovic, Armin Ensser, Alexander S. Hahn

## Abstract

Kaposi’s sarcoma-associated herpesvirus (KSHV) is a human tumor virus. It is associated with Kaposi’s sarcoma, primary effusion lymphoma, and multicentric Castleman’s disease. KSHV is known to interact with several different receptors, among them heparan sulfate proteoglycans, Eph family receptors, and integrins. We mutated the closely related rhesus monkey rhadinovirus in the known receptor interaction sites for Eph family and Plexin domain containing proteins and found it to still replicate on certain cells. This lytic virus was then used as a selection agent in a genome-wide CRISPR knockout screen, which identified TIM1 and NRP1 as host dependency factors. NRP1 is also host factor for the related Epstein-Barr virus and was recently reported to promote KSHV infection, which we confirm even if it functions with low efficiency on most cells and became functional only after ablation of the Eph receptor interaction. Further analysis through overexpression demonstrated that Tim-1 and the related Tim-4 are strong mediators of RRV and KSHV infection, in particular in the absence of other receptor interactions and even more pronounced for a KSHV mutant deleted in glycoprotein K8.1. Both Tim-1 and Tim-4 are heavily O-glycosylated phosphatidylserine (PS) receptors. For KSHV in particular, experiments with mutated Tim-1 and comparison to Ebola virus glycoprotein-driven entry indicate that the interaction with Tim-1 occurs through PS-binding by Tim-1 and suggest additional interaction in a PS-independent manner. The mucin-like domain of Tim-1 is required for optimal receptor function. The use of Tim proteins for entry is a novelty for herpesviruses and underscores the unique biology of KSHV and RRV.

## INTRODUCTION

KSHV is associated with a significant disease burden, in particular in Sub-Saharan Africa where Kaposi’s sarcoma ranks among the most frequent types of cancer^1,2^. In addition, KSHV is associated with primary effusion lymphoma, multicentric Castleman’s disease, and KSHV-positive large B cell lymphoma, as well as potentially with osteosarcoma^1,3,4^, and with an inflammatory cytokine syndrome^5^. KSHV infection of host cells occurs through interaction of the virus with a number of cellular factors, among them heparan sulfate proteoglycans^6–8^, integrins^9,10^, and Eph family receptor tyrosine kinases^11–14^ (reviewed in ^15^). These interactions are generally thought to result in endocytosis of the viral particle and then membrane fusion from endocytic compartments, as various endocytic routes have been found to be important for productive entry of KSHV and it is sensitive to inhibition of the vacuolar ATPase^14,16–18^. For both KSHV and the related rhesus monkey rhadinovirus (RRV) experiments aimed at ablating certain receptor interactions, such as those with integrins^16,19^ or Eph family receptors^11,14,20–22^, mutating receptor interaction sites^12,21^ or deleting individual glycoproteins^23–25^ did not result in complete loss of infectivity, which suggests the existence of additional cellular factors that mediate entry. KSHV and RRV exhibit high similarity with regard to cell tropism, associated disease spectrum and in part receptor usage as both viruses use Eph family receptors^12,14,22^. Interestingly, studies with RRV, that does not interact with integrins or at least not in a similar way as KSHV does^10^, have shown that RRV even despite ablation of all known receptor interactions of the gH/gL glycoprotein complex (RRV^gHΔ^^21–27/AELAAN^, from here on named RRVdM) remains replication competent^22^. Unlike KSHV, RRV is a highly lytic virus on many cells. We therefore decided to use the RRVdM virus in combination with a genome-wide CRISPR library to screen for dependency factors that allow RRVdM to replicate in the absence of known receptor interactions.

## RESULTS

### Genome-wide CRISPR knockout screen identifies TIM1 and NRP1 as potential host dependency factors for RRV

After transduction of the Brunello genome-wide CRISPR library^26^ into A549 cells, the transduced cells were infected with RRVdM or RRV wt (both in an RRV-YFP reporter virus background) to select for cells that are resistant to these viruses (Fig. 1a). A549 were chosen for RRVdM’s ability to replicate on these cells. After no more cytopathic effect was visible, the surviving cells were harvested and the sequences of the transduced sgRNAs in these cells were retrieved. Among the top 50 enriched genes in the surviving cell population we found sgRNAs targeting EphB2 in the population selected for with RRV wt (Fig. 1 b). EphB2 out of the 14 Eph family receptors in humans is the Eph with the second highest affinity for the RRV gH/gL complex and according to the human protein atlas between the three RRV-binding receptors EphB1, EphB2, and EphB3 the receptor with by far the highest level of expression in A549 cells^14,27,28^. Therefore, we were confident that our screen is able to identify host factors. In the cells selected with RRVdM, we identified a number of membrane proteins among the top 50 enriched target genes using FunRich^29^, for which we designed a maximum of four additional sgRNAs. After testing those small guide RNAs in a re-screen with RRV wt (Fig. 1c), those targeting the HAVCR1 (hepatitis A virus cellular receptor 1) gene, also called TIM1 and coding for the Tim-1 protein (Uniprot Q96D42, also called T-cell immunoglobulin mucin receptor 1) came out on top. There are three related TIM genes in humans, TIM1, TIM3, and TIM4. The Tim-1 protein binds phosphatidylserine (PS), thereby for example binding to apoptotic cells^30^. Tim-1 also bears a high molecular weight mucin-like glycosylation on its mucin-like domain, that is of different lengths in Tim-1, Tim-3, and Tim-4 and that is anchored to the membrane by a short stretch of amino acids and a transmembrane domain followed by a small intracellular part^31^. Interestingly, TIM1 and the related TIM4 are known to augment infection by a variety of viruses^32,33,33–40^. Hepatitis C virus, arboviruses like Tick-borne encephalitis virus, Dengue, West Nile and Japanese encephalitis virus, Chikungunya virus, and filoviruses like Lassa and Ebola virus (EBOV) but so far no herpesviruses have been described to use Tim-1 for cell entry. We therefore analysed TIM1 further.

**Fig. 1:**
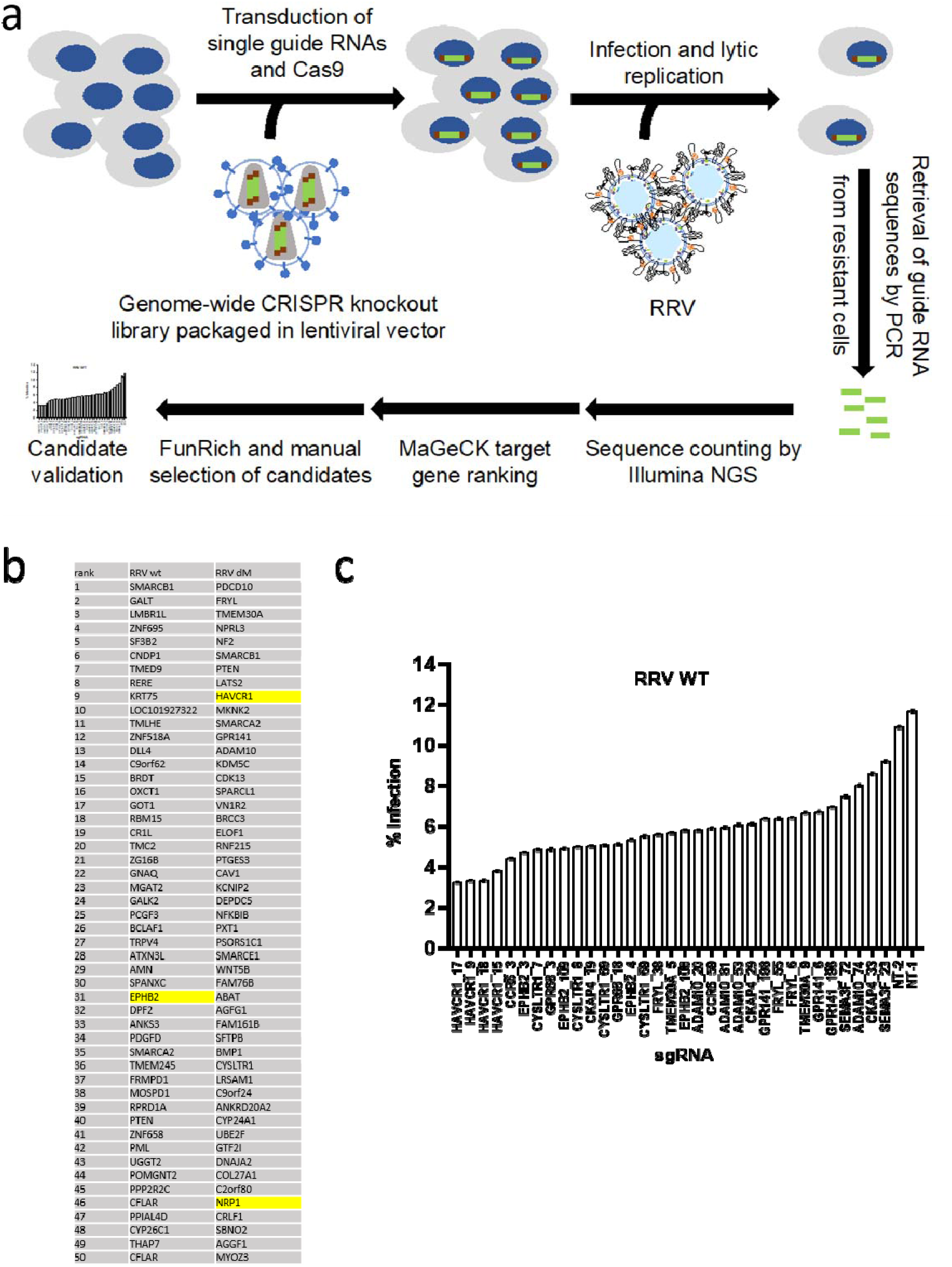
CRISPR-based screen for RRV host factors. (a) Schematic representation of the screening workflow. (b) List of top 50 enriched genes after selection compared to the transduced library without infection. EPHB2, HAVCR1 (TIM1) and NRP1 are highlighted. (c) Rescreening of selected candidate genes with independent sgRNAs.

### Knockout or knockdown of TIM1 reduces infection by RRV and KSHV

To test function of TIM1 in the infection of different cells we used CRISPR-mediated knockout of TIM1. Testing a number of adherent cell lines, we found knockout of TIM1 to significantly reduce infection by KSHV and RRV in SLK and A549 cells (Fig. 2a&b), which both express the Tim-1 protein as assayed by cell surface staining and Western blot (Fig. 2a&b, middle and rightmost panels). It should be noted that human foreskin fibroblasts (HFF) and human ubilical vein endothelial cells (HUVEC) did not express detectable levels of Tim-1 as assayed by Western blot, and consequently knockout/knockdown of TIM1 did not affect infection (Fig. 2c&d), affirming the specificity of the observed effects and ruling out off-target activity of the CRSIPR/Cas9 system against KSHV or RRV. Interestingly, there was an obvious discrepancy between Tim-1 expression as assayed by Western blot or surface expression on A459 cells (Fig. 2b). While knockout/knockdown was clearly visible in Western blot, surface expression was unchanged upon knockdown, and the signal was only threefold over background. This suggests that most of the Tim-1 protein resides intracellularly, probably even all of it, if the weak positivity by flow cytometry represents background rather than true signal.

**Fig. 2:**
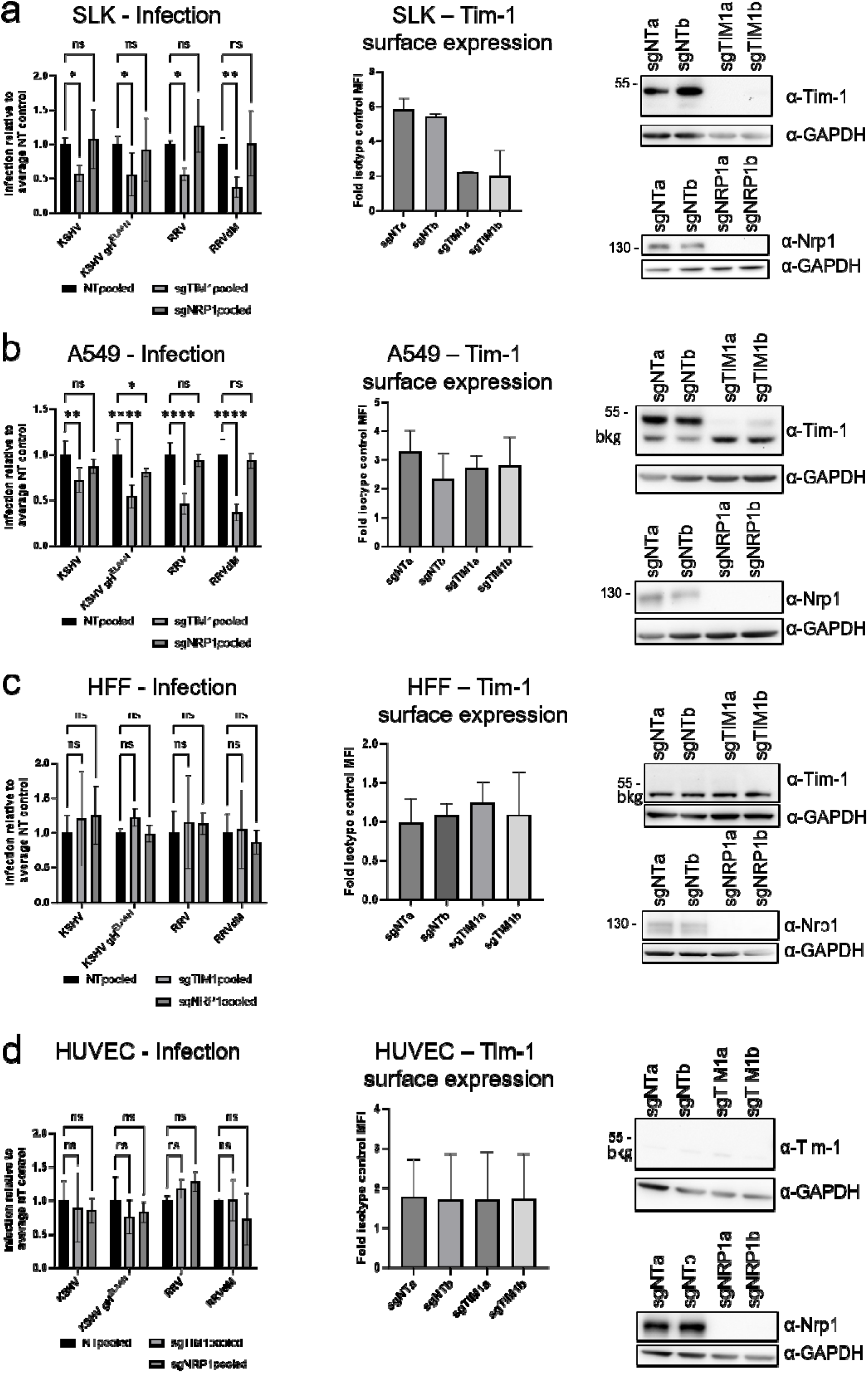
Reduction of KSHV and RRV infection by TIM1 knockout on Tim-1-positive but not on Tim-1-negative cells. Tim-1-positive (a) SLK cells and (b) A549 or Tim-1-negative (c) HFF and (d) HUVEC were transduced with Cas9 and two different non-targeting sgRNAs (sgNT) or two different sgRNAs targeting TIM1 or NRP1. The experiment was performed in biological triplicates for each sgRNA construct on two (a, only one NRP1-sgRNA in one experiment) or three (b, only two for NRP1; c only two for KSHVgH^ELAAN^ and RRVdM) different occasions, results for different sgNTs and different TIM1-targeting sgRNAs were pooled. ns non-significant, * p<0.05, ** p<0.01, **** p<0.0001, two-way ANOVA, Dunnett‘s correction for multiple comparisons. Surface staining and flow cytometry analysis (middle panel) and Western blot analysis (right panel) to confirm knockout/knockdown of Tim-1 and Nrp1 protein levels was performed.

### An antibody to Tim-1 blocks KSHV infection of Tim-1-positive cells

To further ascertain the specificity of the effects observed upon Tim-1 knockout, we targeted Tim-1 with a well-characterized antibody^41^. Pre-treatment of cells with Tim-1 antibody reduced infection of SLK and A549 cells significantly. We achieved 63% (A549) to 70% (SLK) inhibition (Fig. 3) at the highest concentration, where the effect was in saturation (not shown). Infection of HUVEC and HFF, which do not express Tim-1 to detectable levels, was not significantly inhibited.

**Fig. 3:**
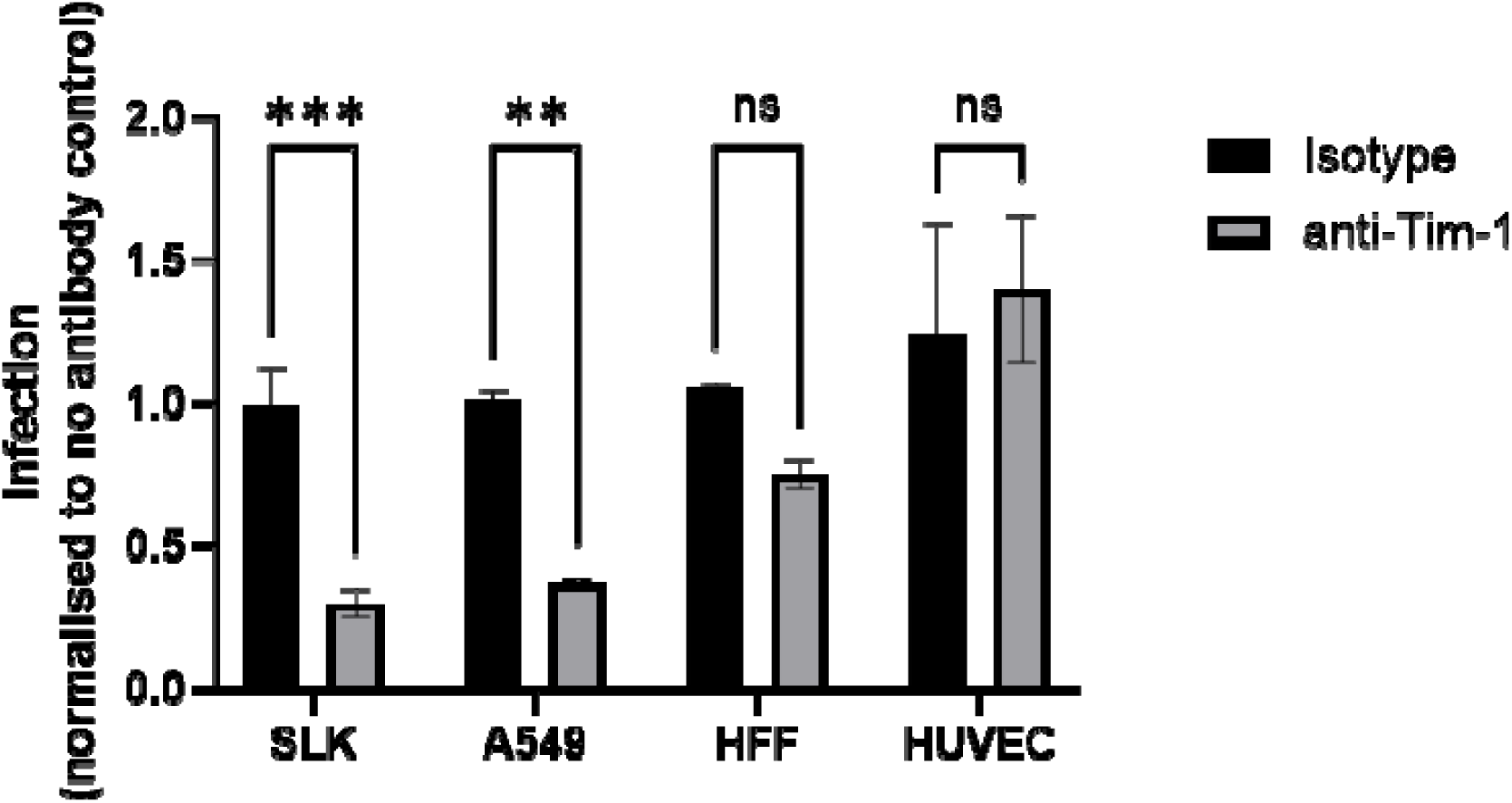
A monoclonal antibody to Tim-1 inhibits KSHV infection of Tim-1-positive SLK and A549 cells, but nor of Tim-1-negative HFF and HUVEC. The cells were pre-incubated with the indicated antibodies for 30 min prior to addition of KSHV. Final antibody concentration was 2.5 µg/ml. ** p<0.01, *** p<0.001, two-way ANOVA, Sidak‘s correction for multiple comparisons.

### Knockout or knockdown of NRP1 slightly reduces infection by KSHV in a cell-specific manner and when the Eph-interaction is perturbed

Neuropilin 1 (Nrp1) is a known receptor for the closely related EBV^42^, and it was recently published to also promote infection of mesenchymal stem cells by KSHV through interaction with glycoprotein B (gB)^43^. We therefore limited our studies addressing Nrp1 function to verifying its role during KSHV and RRV infection in a limited set of cells. Here, we found that reducing expression of NRP1 slightly reduced infection of A549 by KSHV and RRV, albeit to a lesser extent than was observed for TIM1, and the reduction in infection did not reach significance except for infection with the KSHV gH^ELAAN^ mutant on A549 cells (Fig. 2b), suggesting that NRP1 becomes more important when Eph family receptors are not available. It should be noted that while knockout of TIM1 in A549 was not entirely complete, knockout of NRP1 resulted in the complete absence of detectable expression (Fig. 2b, rightmost panel). Therefore, NRP1 function seemed to be cell-specific and did not obviously correlate with its expression levels as knockout/knockdown did not decrease infection of HFF and HUVEC cells, despite appreciable expression levels (Fig. 2).

### Overexpression of TIM1 and TIM4 strongly enhances KSHV and RRV infection

293T cells are essentially negative for TIM1 expression (Human Protein Atlas, proteinatlas.org)^27,28^. We therefore decided to use these cells for gain-of-function experiments and recombinantly expressed Tim-1 or the related Tim-3 and Tim-4 by means of lentiviral transduction on these cells. Tim-1 and Tim-4 are highly similar, whereas Tim-3 is more dissimilar and almost completely lacks the large mucin-like, O-glycosylated domain of the other two TIMs^31^. Overexpression of Tim-1 or Tim-4 increased infection by RRV and KSHV significantly (Fig. 4a&c), whereas Tim-3 expression only resulted in modest enhancement, if at all, with both wt KSHV and RRV as well as mutants of both viruses mutated in the Eph interaction interface (Fig. 4b&d). Surprisingly, infection by KSHVΔK8.1 was enhanced to practically the same degree through overexpression of Tim-3 in this assay (Fig. 4e), suggesting that also Tim-3 can meaningfully augment KSHV infection, but only under circumstances where e.g. glycoprotein K8.1 is not functional.

**Fig. 4:**
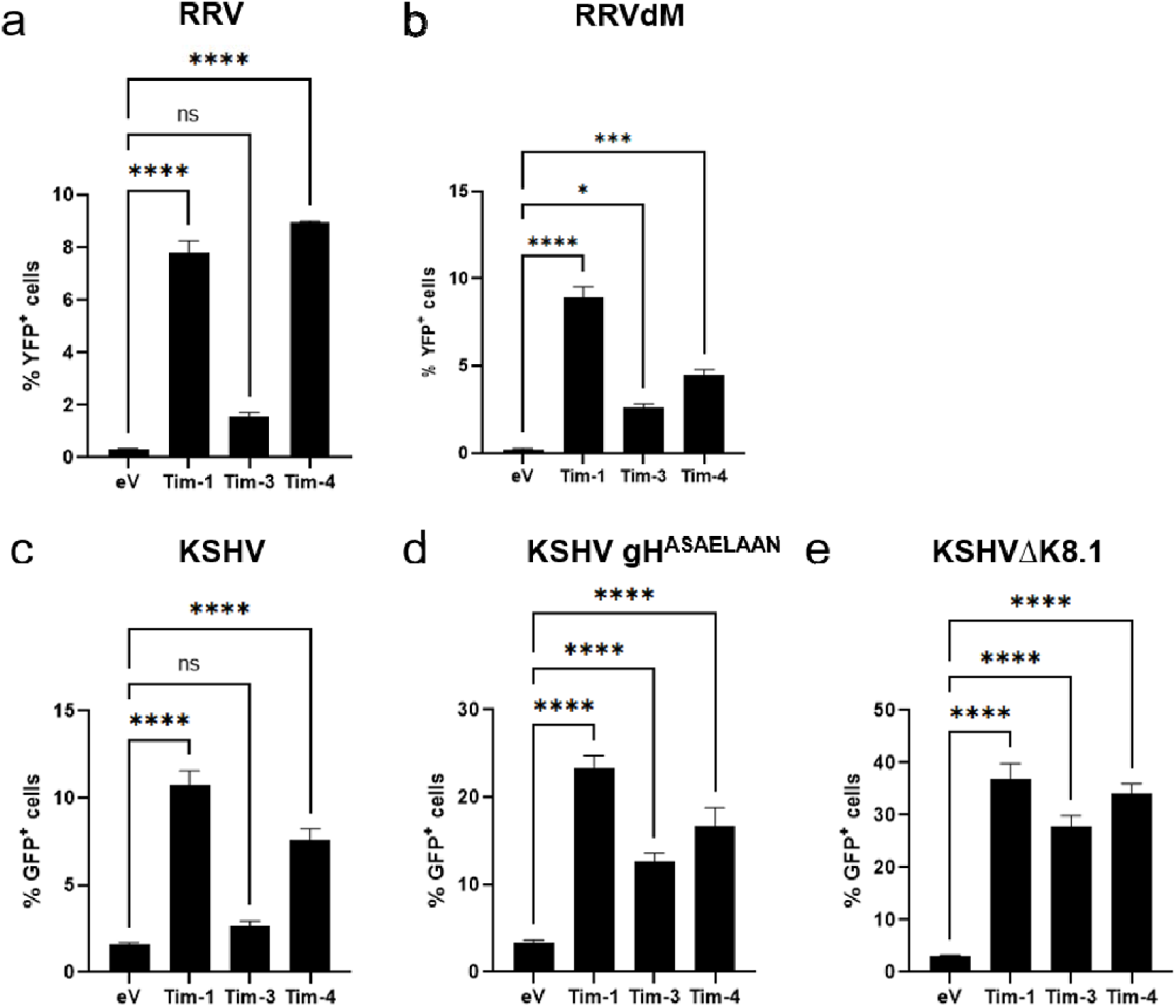
Directed expression of Tim-1, Tim-3, and Tim-4 enhances infection by KSHV and RRV. 293T cells were transduced with cDNAs encoding transcripts for Tim-1, Tim-3, or Tim-4 under the control of the strong CMVie promoter. After transduction and brief selection of transduced cells, the resulting cell pools were infected with RRV (a), RRVdM (b), KSHV (c), or the indicated mutant KSHV gH^ASAELAAN^ (d) or KSHVΔK8.1 (e). The experiment was performed in triplicate. ns non significant, * p<0.05, ** p<0.01, *** p>0.001, **** p<0.0001, two-way ANOVA with Dunnett‘s correction for multiple comparisons.

### Soluble Tim-1 blocks infection by KSHV and RRV and interacts with KSHV through PS and through PS-independent mechanisms

In order to probe whether cellular Tim-1 modulates KSHV/RRV infection indirectly or can be competed by a soluble version of the receptor, we preincubated KSHV or RRV with soluble Tim-1-Fc as a decoy receptor. This resulted in significant, dose-dependent inhibition of KSHV and RRV on the Tim-1-positive cell line A549 (Fig. 5a).

**Fig. 5:**
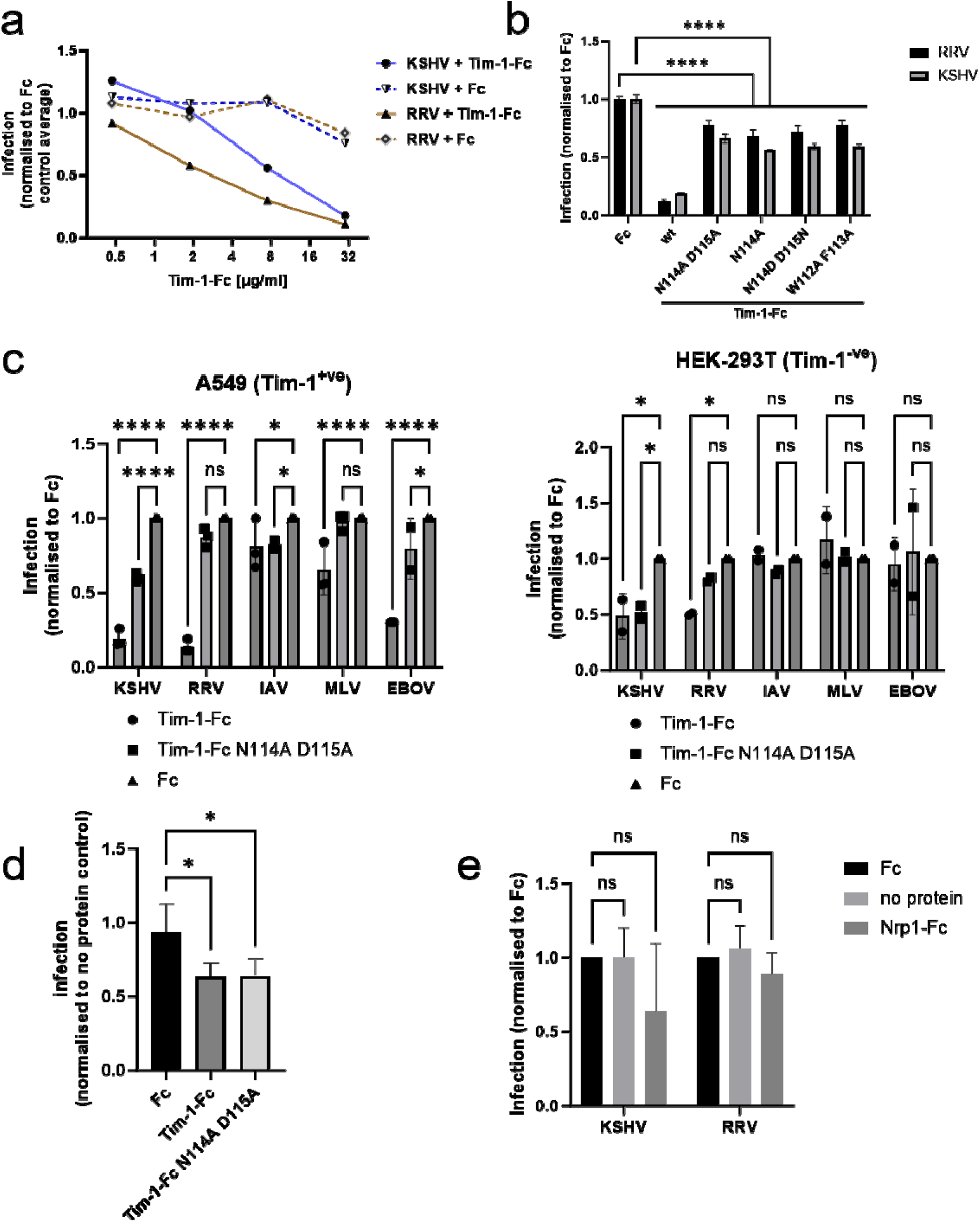
Soluble Tim-1 blocks infection in a PS-dependent and PS-independent fashion. (a) KSHV or RRV were pre-incubated with soluble Tim-1-Fc fusion protein prior to infection of A549 cells. Values represent the average of triplicate samples, error bars were omitted for clarity. (b) KSHV or RRV were pre-incubated with soluble wt Tim-1-Fc fusion protein or Tim-1-Fc bearing the indicated mutations. **** p<0.0001, two-way ANOVA, Dunnett‘s correction for multiple comparisons. (c) Tim-1-positive A549 or Tim1-negative 293T cells were infected with KSHV, RRV or lentiviral pseudotypes carrying the indicated viral glycoprotein (IAV = infuenza A virus hemagglutinin and neuraminidase, MLV = murine leukemia virus envelope protein, EBOV = Ebola virus glycoprotein). ns non-significan, * p<0.05, **p<0.01, ***p<0.001, **** p<0.0001. D) 293T cells were infected with KSHV gH mutant virus that does not use Eph family receptors and that was pre-incubated with mutant Tim-1-Fc or Tim-1-Fc N114A D115A at 30µg/ml. * p<0.05, one-way ANOVA, no correction for multiple comparisons. (e) A459 cells were infected with KSHV or RRV preparations that were pre-incubated with soluble Nrp1-Fc or Fc protein at 35 µg/ml. The experiment was performed twice in triplicates with two independent protein preparations. ns non-significant, Dunnett‘s correction for multiple comparisons.

We next asked whether the inhibitory effect of soluble Tim-1-Fc was dependent on PS binding. We compared a number of different soluble receptor proteins that were mutated in the PS-binding site (Fig. 5b). None of the PS-binding site mutations fully abrogated inhibition of infection by these soluble proteins.

To further investigate this observation, we compared inhibition of KSHV, of RRV and of lentiviral pseudoviruses (LP) bearing either influenza hemagglutinin and neuraminidase (IAV LP), murine leukemia virus envelope protein (MLV LP), or EBOV glycoprotein (EBOV LP) (Fig. 5c). On Tim-1-positive A549 (Fig. 5c, left panel), we observed significant inhibition of all viruses. While inhibition of IAV LP and MLV LP did not exceed 20% for IAV LP and 40% for MLV LP and EBOV LP was inhibited by approx. 70%, inhibition of KSHV and RRV reached approx. 80%. Interestingly, KSHV and to a lesser degree also EBOV LP and IAV LP but not RRV were still significantly inhibited by soluble Tim-1 N114A D115A, which is mutated in the PS binding pocket, specifically where the charged head group of PS binds^44,45^, which is in line with reports of PS-independent interactions of EBOV GP with Tim-1^41^. On 293T cells, which are negative for Tim-1, none of the pseudotyped lentiviruses was inhibited by soluble Tim-1-Fc (Fig. 5c, right panel). RRV was inhibited by wt soluble Tim-1-Fc. KSHV on the other hand was inhibited by both soluble versions of wt and PS-binding pocket-mutated Tim-1-Fc, and to the same extent. This inhibition of KSHV by PS-binding-deficient Tim-1 decoy receptor on Tim-1-negative cells could be interpreted that either Tim-1 binds to KSHV particles, or that Tim-1 binds to another cellular receptor of KSHV and thereby blocks entry. This is a valid hypothesis as Tim-1 and Tim-4 reportedly interact with EphA2^46^. Even so, a KSHV gH ASAELAAN mutant, that does not use EphA2 for entry, was also potently inhibited by soluble Tim-1-Fc and PS-binding-mutated Tim-1-mut-Fc (Fig. 5d). This suggests that blocking of EphA2 by soluble Tim-1 is not a relevant mechanism.

### Soluble Neuropilin 1 exhibits a trend toward inhibition of KSHV and RRV infection

We also tested inhibition of KSHV and RRV by soluble Nrp1-Fc decoy receptor at 35 µg/ml (Fig. 5e). We observed a trend towards inhibition of KSHV or RRV infection of A549 cells. The effect was considerably more pronounced for KSHV than for RRV. The comparatively high dose that was needed for an effect may be rooted in the fact that Nrp1 binding to furin-processed proteins like KSHV gB is of low affinity^43^. This requirement for a high dose and an effect potentially not yet in saturation may also have contributed to the high variance, preventing statistical significance from two repeats with different protein preparations, which may have slightly differed in their biological activity. This is a factor we cannot fully control for. This problem may be overcome by performing additional repeat experiments, but we deemed this unnecessary as our results in principle confirm those by Lu et al.^43^, and NRP1 was not the focus of our study.

### Mutations in the phosphatidylserine-binding pocket of Tim-1 reduce but do not abrogate receptor function for KSHV and RRV

To further analyse a potential PS-independent function of Tim-1 during KSHV infection we transduced 293T cells with expression constructs coding for wt Tim-1 or Tim-1 mutated in the PS-binding site, either in the described binding pocket for the head group (N114, D115) or in the hydrophobic interaction site (W112, F113) that supposedly binds to the lipid part of PS (Fig. 6). Only enhancement by full-length Tim-1 reached significance in this set of three averaged experiments for KSHV, KSHV gH^ASAELAAN^ (Fig.6a) or RRV and RRVdM (Fig. 6b). All mutations decreased enhancement of RRV and KSHV infection but the PS-binding site mutations did not fully abrogate it, even if enhancement by these mutants did not reach significance, which was owed to the number of comparisons and the high variance as the data represents three independent sets of experiments on different occasions in triplicates, which increases variability. In a single experiment in triplicates performed in parallel with less conditions, PS binding pocket-mutated Tim-1 clearly enhanced infection of KSHV wt and more so of KSHVΔK8.1 significantly also in a statistical sense (Fig. 6c), compatible with the hypothesis of PS-dependent and PS-independent functions contributing to infection. Noticeably, KSHVΔK8.1 was enhanced to a considerably higher degree than wt KSHV.

**Fig. 6:**
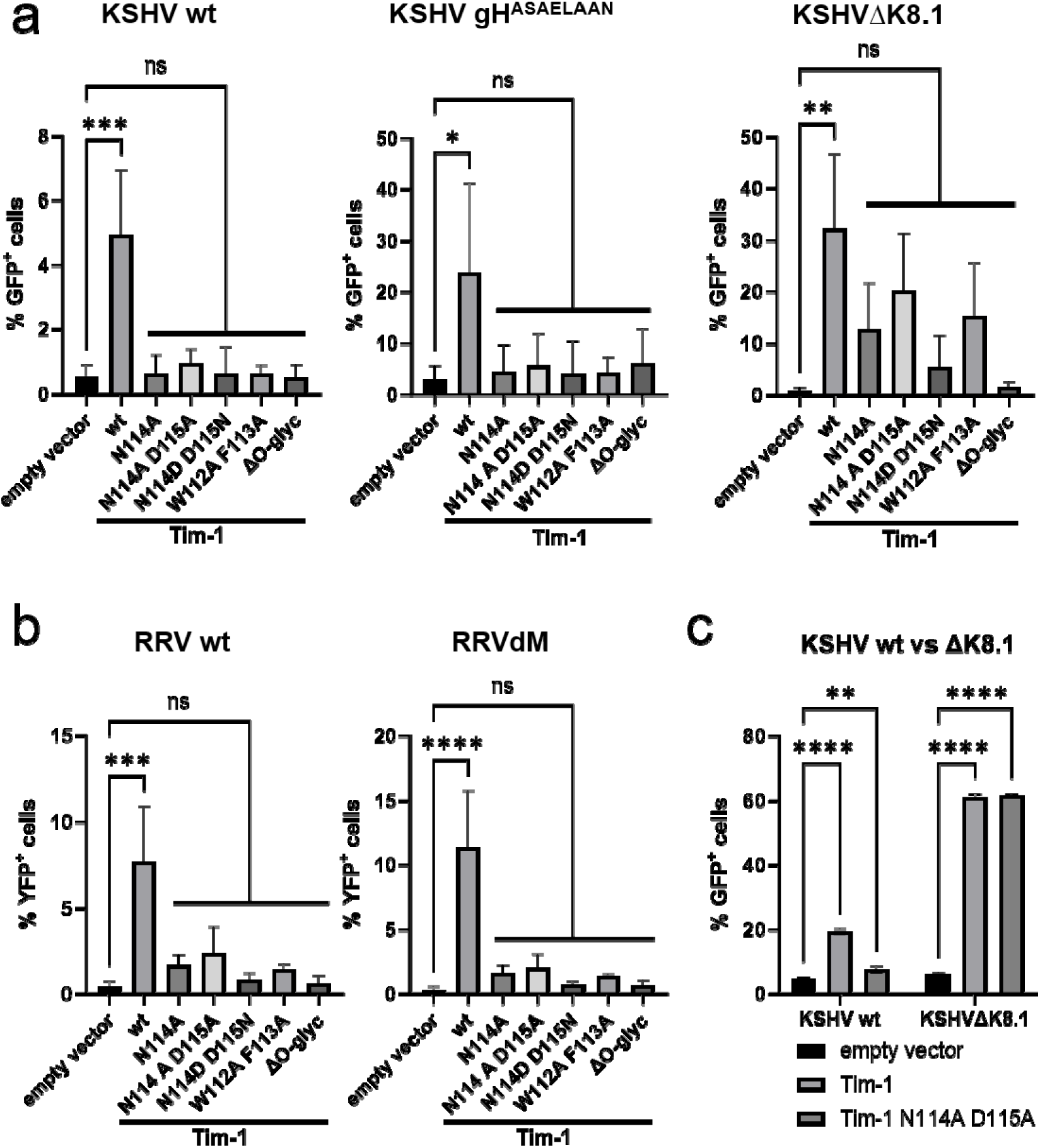
The phosphatidyl serine-binding site of Tim-1 is important but potentially not essential for Tim-1 receptor function. 293T cells were transduced with cDNAs encoding Tim-1 or the indicated Tim-1 mutants under the control of the strong CMVie promoter. After transduction and brief selection of transduced cells, the resulting cell pools were infected with (a) KSHV wt and the indicated mutant viruses or (b) RRV wt or the indicated mutant viruses. * p<0.05, ** p<0.01, *** p>0.001, **** p<0.0001, one-way ANOVA with Dunnett‘s correction for multiple comparisons. c) Separate experiment to compare Tim-1 and Tim-1 N114A D115A with regard to function in KSHV infection. ns non-significan, * p<0.05, ** p<0.01, *** p>0.001, **** p<0.0001, two-way ANOVA with Dunnett‘s correction for multiple comparisons.

### Regions outside the phosphatidylserine-binding pocket contribute to Tim-1 receptor function for KSHV infection

Interestingly, a mutant that is deleted in the O-glycosylation-rich mucin domain (Tim-1ΔO-glyc) did not enhance infection by KSHVΔK8.1 or RRV (Fig. 7a&b). KSHVΔK8.1 was chosen as it was the virus that responded to expression of Tim-1 or Tim-1 mutants with the most pronounced increase in infection. In order to control for sufficient surface expression of this Tim-1ΔO-glyc mutant, we constructed a variant with an N-terminal Flag-tag inserted right after the signal peptide. In this system, Tim-1ΔO-glyc was expressed on the cell surface at a level that was considerably lower compared to that of full length Tim-1 (Fig. 7c), but it was clearly detectable. In immunofluorescence analysis, a larger fraction of the mutant protein lacking the O-glycosylated stalk seemed to localise intracellularly (not shown). The pronounced differences in expression levels preclude a definitive conclusion on the role of the O-glycosylated stalk.

**Fig. 7:**
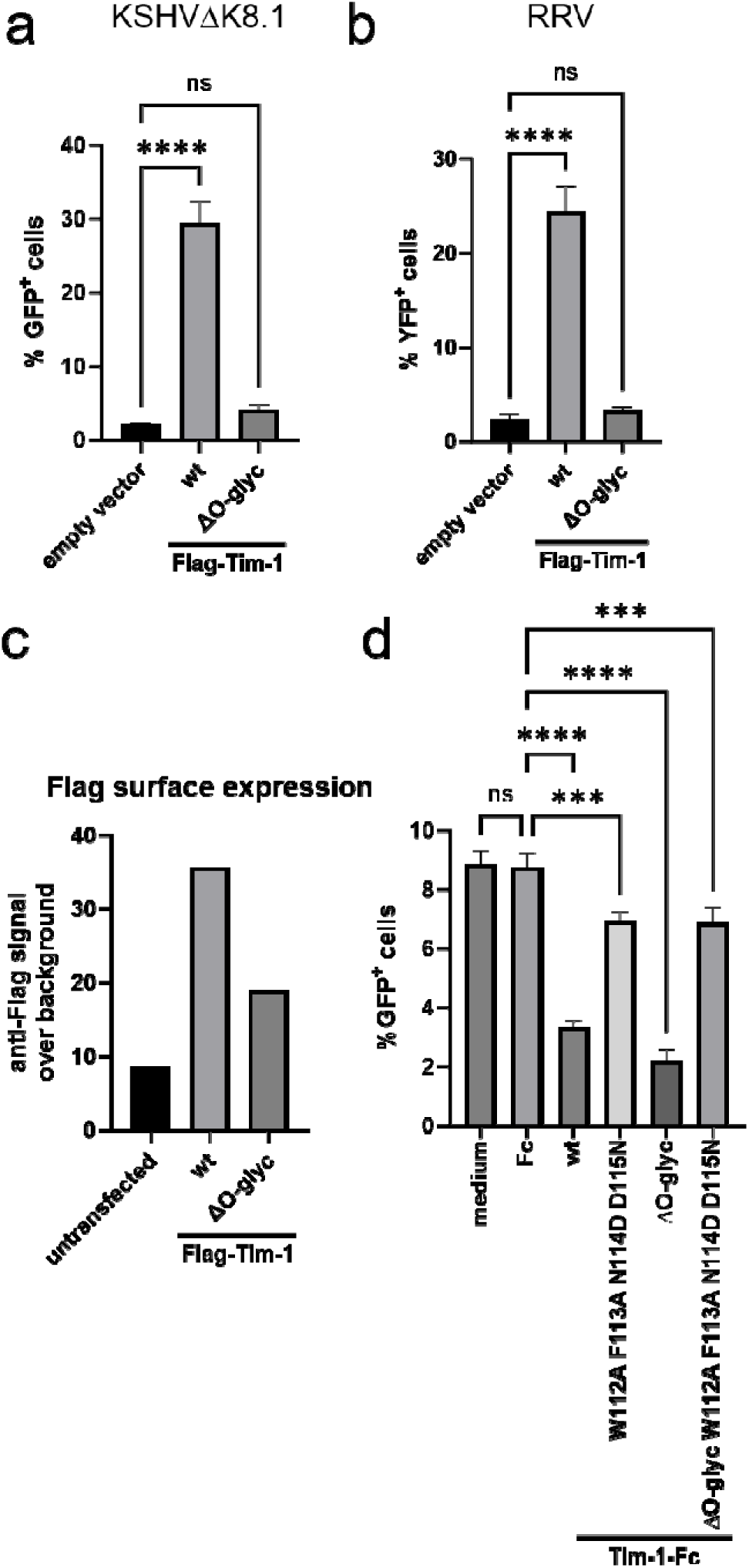
Evidence that multiple regions of Tim-1 function during KSHV infection through direct interaction or indirect effects. 293T cells were transduced with lentiviral vectors encoding Tim-1 or Tim1-ΔO-glyc, both with an N-terminal Flag-tag. The cells were then infected with (a) KSHV or (b) RRV. Flag surface expression was analysed by flow cytometry. (d) Inhibition of KSHV infection by soluble Tim-1-Fc or soluble Tim-1-Fc bearing the indicated mutations, all at 30 µg/ml. * p<0.05, ** p<0.01, *** p>0.001, **** p<0.0001, ordinary one-way ANOVA with Dunnett‘s correction for multiple comparisons.

A soluble version of Tim-1ΔO-glyc on the other hand inhibited KSHV (Fig. 7d) as did versions of Tim-1-Fc that were mutated at 4 different residues of the PS-binding pocket (W112A F113A N114D D115N), essentially completely destroying this motif. Concomitant deletion of the O-glycosylated domain and destruction of the PS-binding motif in the soluble Tim-1-Fc construct (ΔOGlyc W112A F113A N114D D115N) still did not fully abrogate inhibition of KSHV by this molecule. Overall, this data indicates that the mucin-like domain is critical for either receptor function or correct expression or localisation of Tim-1 or all three but not for inhibition by Tim-1 as a soluble decoy receptor. This clearly implicates the immunoglobulin-like, PS-binding domain of Tim-1 in the interaction with KSHV, and according to our data both in a PS-dependent and PS-independent manner, whereas the O-glycosylated stalk may play a different functional role.

### TIM1 interacts with KSHV at a post attachment step

Given TIM1’s ability to bind PS in viral membranes, attachment would be a plausible step where TIM1 functions during infection. However, we found that soluble Tim1-Fc did not decrease attachment at a concentration that decreased infection (Fig. 8a). Heparin on the other hand clearly decreased attachment. We therefore further investigated the temporal association of Tim-1 with viral particles after initial binding. We found a considerable part of Tim-1 distributed intracellularly in SLK cells. Association of Tim-1 with KSHV particles after attachment at 4°C as assayed by colocalisation with fluorescently tagged KSHV virions was visible, with approx. 10% of the KSHV capsid signal colocalized (Fig. 8b&c middle panel). After incubating the attached virus further at 37°C, colocalisation of KSHV capsids with Tim-1 increased significantly but at this point seemingly occurring more in intracellular areas of accumulated Tim-1 (Fig. 8b&c right panel). Overall, compatible with the results of the attachment assay (Fig. 8a), Tim-1 did not interact extensively with KSHV directly upon binding of the virus, but colocalisation increased over time, implicating a role post attachment.

**Fig. 8:**
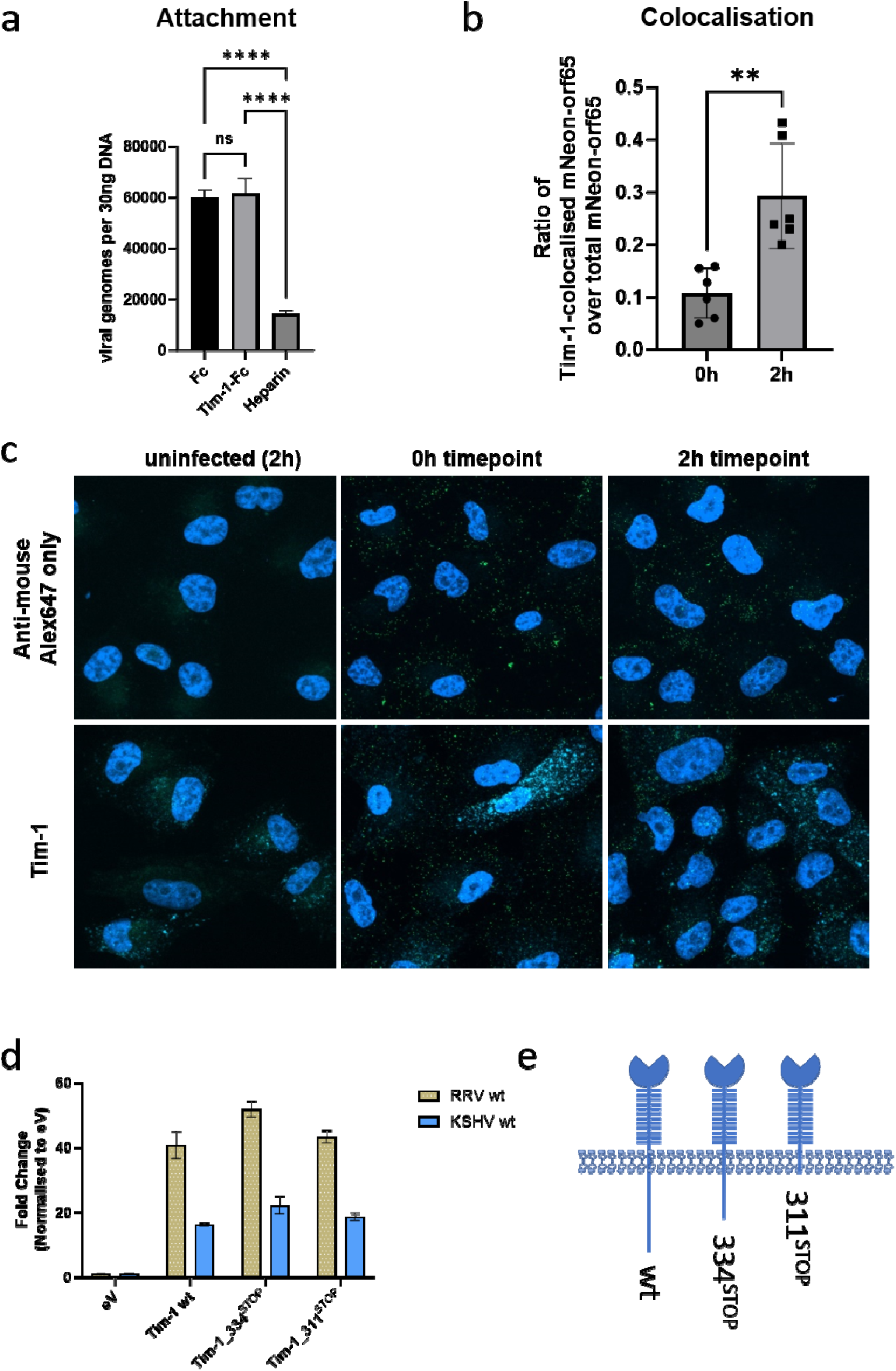
Tim-1 functions at a post-attachment step without an obvious role for the intracellular domain. a) Attachment of KSHV to SLK target cells was measured after preincubation of the virus with Tim-1-Fc or Fc at 10µg/ml, or as a positive control with Heparin at a concentration of 1250 U/ml. **** p<0.0001, one-way ANOVA with Dunnett‘s correction for multiple comparisons. b) SLK cells were incubated with KSHV-mNeonGreen-orf65 at 4°C and then washed in ice-cold PBS. The cells were then either fixed immediately or incubated for another 2h at 37°C before fixation for immunofluoscence staining for Tim-1 and analysis by confocal microscopy. Colocalisation of KSHV capsids with Tim-1 was assessed for both conditions. ** p<0.01, t-test for unpaired samples. c) Representative images as used for analysis in (B). (d) Intracellular truncation variants of Tim-1 were recombinantly expressed in 293T cells by means of lentiviral transduction, the cells were infected with KSHV or RRV, and infection was quantified by flow cytometry. (e) Schmatic representation of truncation mutants used in (d).

### The intracellular part of Tim-1 is dispensable for receptor function

As the association of KSHV with Tim-1 increased during the first 2h of infection but did not contribute to attachment of KSHV, one hypothesis would be that Tim-1 augments infection through directly recruiting intracellular factors that foster endocytosis or macropinocytosis through its intracellular domain. To address this question, we overexpressed Tim-1 with truncating deletions that remove approximately half or the complete intracellular part. Surprisingly, these intracellularly truncated Tim-1 molecules performed equally well as receptors for KSHV and RRV (Fig. 8d&e), at least within the limits of our experimental system.

## DISCUSSION

Our findings demonstrate that Tim-1 and the related Tim-4 can serve as receptors for KSHV and the related RRV, while Tim-3 is considerable less efficient at promoting infection. We use the term receptor here because: i) TIM1 allows KSHV/RRV to efficiently infect cells without contributing appreciably to virus attachment and colocalisation increased over time after binding. ii) Soluble Tim1-Fc inhibits KSHV/RRV infection, as to be expected from the soluble decoy version of a cellular receptor. Similar to other molecules termed receptors for KSHV and RRV^9,11,47^, TIM1 and TIM4 are not essential for these two viruses. We made similar findings for NRP1, but with the caveat that we were unable to demonstrate a direct interaction of Nrp1 with viral glycoproteins (not shown), which is likely rooted in the choice of our methods. Lu et al. demonstrated interaction of Nrp1 with gB by GST pulldown and biolayer interferometry, which can detect more transient interactions with lower affinities. Nrp1 affinity for gB was reported as 263nM, which is by over one order of magnitude lower than what was measured for the gH/gL-EphA2 interaction, which is below 10nM^13,48^. Even so, we clearly observed inhibition by soluble Nrp1-Fc, indirectly confirming its interaction with the viral particle, likely through gB as published by Lu et al.

Findings with KSHV and RRV so far raise the question whether there is a single one essential receptor for KSHV and RRV. The broad host and cell tropism in vitro and the ability to infect in the absence of known receptor interactions as reported by e.g. TerBush et al.^19^ argue against this hypothesis. Rather, the current state of knowledge suggests that the known receptors may act synergistically to allow the virus access to a compartment where membrane fusion occurs, or may foster virus binding as observed in other studies for integrins and Nrp1^8,43^. Reports on a specific fusion receptor, xCT, remain conflicting^49,50^, and roles in attachment, endocytosis, and fusion for individual host receptors may overlap and not be exclusive, as is the case for the Eph family of receptors for KSHV and RRV^12,21,22,47^, which can trigger both fusion and endocytosis^11,51,52^. In line with this notion, we initially identified the Tim-1 encoding HAVCR1 gene as critical for RRV entry using a virus mutant deleted for the known receptor interactions with Eph and Plxdc family proteins as selection agent, whereas our screen with RRV wt reported the known EPHB2 receptor gene among the 50 highest ranking genes, but not TIM1 or NRP1 (Fig. 1b). Likewise, observed effect sizes upon modulation of TIM1 expression were slightly larger when virus mutants negative for other known receptor interactions or deleted in the glycoprotein K8.1 gene locus were used for infection experiments (Fig. 4d&e).

While for e.g. the EBOV GP-pseudotyped lentivirus the interaction with Tim-1 seemed to be almost entirely dependent on binding of Tim-1 to PS, as e.g. soluble Tim-1 mutated in the PS binding domain had only a very limited effect on infection, the interaction of Tim-1 with RRV and in particular KSHV appears to be more complex. There clearly is a PS-dependent component as evidenced by strongly decreased receptor function upon mutation of the known PS-binding pocket (Fig. 6), but there also is strong evidence for a PS-independent interaction of Tim-1 with KSHV as evidenced by inhibition of infection of Tim-1-negative 293T cells by soluble Tim-1-Fc fusion protein mutated in the PS binding pocket (Fig. 5c) and by enhancement of infection by overexpression of such Tim-1 mutants, particularly obvious when a K8.1 deletion mutant of KSHV was used (Fig. 6). Interaction of PS-binding-negative Tim-1 mutant proteins with KSHV and with the related RRV differed in that regard to some degree. Results with RRV were more similar to EBOV GP pseudotyped particles in that e.g. a soluble Tim-1 protein with mutated PS-binding pocket did not have an inhibitory effect on EBOV GP-pseudotyped particles on Tim-1-negative 293T cells, unlike what was observed for KSHV, which was inhibited (Fig. 5c). For EBOV a PS-independent Tim-1 interaction has been suggested^41^, but the nature of this interaction seems to differ from the interaction of Tim-1 with KSHV. Inhibition of KSHV by soluble Tim-1 mutated both in the PS-binding site and deleted in the O-glycosylated stalk (Fig. 7d) suggests that KSHV interacts with the N-terminal domain of Tim-1 also outside the PS binding pocket. Our efforts to identify Tim-1-interacting viral glycoproteins so far did not identify a viral binding partner, even if we tested a range of KSHV glycoproteins (not shown). It should be noted that our approach was limited to immunoprecipitation and to cell-surface binding of soluble Tim-1-Fc fusion protein to cells recombinantly expressing viral glycoprotein complexes, analogous to cell surface staining using antibodies. These techniques may require a certain affinity or slow dissociation between the interaction partners that may not be necessary for receptor function. As Tim-1 can already bind with PS or similar molecules in the viral envelope leading to immobilisation on the membrane, a potential second interaction may require only comparatively low affinity to become functional in this context, which may not be easily amenable to analysis by immunoprecipitation or flow cytometry-based binding assays. Alternatively, Tim-1 may bind another cell-derived component of the viral envelope.

Mechanistically, it is noteworthy that KSHV and RRV, similar to e.g. EBOV, enter via an endocytotic route, in many cases macropinocytosis^14,17,18,51,53^. Tim-1 e.g. on A549 cells seemed to localise mostly to the interior of the cell, as can be deduced from the fact that surface expression as measured by flow cytometry was low or absent and remained unchanged upon TIM1 gene knockout/knockdown, whereas expression as measured by Western blot was visibly reduced or abrogated. We found the colocalization of KSHV with Tim-1 to increase after binding (Fig. 8b&c), compatible with Tim-1 playing a role in the internalisation of the virus rather than during its initial binding to the cell surface, which was unaltered by competition with soluble Tim-1 (Fig. 8a). Our finding that the intracellular domains of Tim-1 were dispensable for receptor function suggest that direct recruitment of intracellular factors by Tim-1 may play at best a minor role. Either, Tim-1 steers KSHV and RRV into regions of the membrane or into vesicles that are particularly conducive to infection or it interacts with the virus surface in a way that facilitates infection, and both functions are not mutually exclusive as demonstrated by the Eph family of receptors. So far, our results indicate that Tim-1 does not have the ability to trigger the fusion machinery of KSHV or RRV directly, unlike e.g. the Eph family of receptors (data not shown).

Unfortunately, different surface expression levels preclude a clear conclusion on the role of the O-glycosylated stalk domain of Tim-1. Nevertheless, our finding that Tim-4, which has a heavily O-glycosylated stalk that is similar to that of Tim-1, functions efficiently as a receptor (Fig. 4), whereas Tim-3, which almost completely lacks a stalk domain, does not, suggests that the stalk domain may fulfil a critical function.

The Tim-1 and Tim-4 interactions fit with the reported tropism of KSHV and RRV. Tim-1 is expressed on T cells, NK cells, dendritic cells and Langerhans cells (Human Protein Atlas, proteinatlas.org)^27,28^, among others, and also broadly on many epithelial cells^41^, while Tim-4 is expressed e.g. on certain T cells, macrophages and plasma cells (Human Protein Atlas, proteinatlas.org)^27,28,54–57^. Closely related viruses of the rhadinovirus RV2 lineage to which RRV belongs were found associated with T cell lymphoma in a report that analysed the association of RV2 viruses with lymphoma in their natural host^58^. Tim-4 is highly expressed on bone marrow plasma cells, and KSHV-EBV co-infected cells exhibit a partial plasma cell phenotype^59^, as do PEL cells with regard to surface markers^60^. Curiously, Tim-1 and Tim-4 are highly expressed on T cells, and Myoung et al. described in 2011 that primary tonsillar T cells are highly susceptible to KSHV infection, in fact much more so than B cells, especially after stimulation, but do not support replication of the virus^61^. The biological consequences of KSHV’s marked tropism for T cells coupled with abortive infection are currently not understood, but our finding of Tim-1 and Tim-4 receptor usage support these previous observations and suggest that there may be an underlying biological rationale that is currently not understood.

A recent study reported a physical association between EphA2 and Tim-4 as well as Tim-1^46^. Evolutionarily, it seems plausible the KSHV and RRV have evolved to use receptors that are in physical proximity to each other. Both EphA2 and the two Tim proteins may localise to areas of the cellular membrane that are particularly suitable for endocytosis of viral particles, which would at least in part explain why EphA2^48,62–66^ and the Tim proteins^34,37,37,40,41,67–69^ are used by a surprising variety of pathogens to promote gain entry to host cells. On the other hand, Tim-1 was clearly functional as a receptor for Eph binding-negative mutants of KSHV and RRV, which indicates that while the Tim-1, Tim-4, and EphA2 may interact and colocalise, their function as entry receptors is not dependent on KSHV/RRV binding to Eph family receptors.

Overall, our findings identify Tim-1 and Tim-4 as novel receptors for KSHV and the related RRV, and we confirm the role of Nrp1 for KSHV and demonstrate its function also in infection by the related RRV, even if the overall contribution of Nrp1 to infection at least in our experiments was limited and even if identified in our initial screen, the reduction in infection upon knockout did not even reach significance in our confirmatory experiments. Soluble Nrp1 effected a trend towards inhibition of KSHV and RRV that did not reach significance, potentially indicating that its binding to gB inhibits similar to inhibition by gB-binding antibodies. Variability in the activity of the two different Nrp1-Fc preparations used by us may have increased variance, which together with limited effect size on the cells used in our assays precluded reaching statistical significance. Even so, overall our results confirm the observations made by Lu et al. but indicate that Nrp1 function may be limited to certain cells, likely those lacking other receptor proteins^43^. The diversity in the receptor interactions of the two rhadinoviruses KSHV and RRV and also of other herpesviruses might either reflect their need to escape the neutralizing antibody response or may enable them to colonise different biological niches. While the Nrp1 interaction with cleaved furin sites of viral surface proteins emerges as a conserved concept across many viruses and also herpesviruses, the interaction with Tim-family proteins by RRV and KSHV is to our knowledge a novum for herpesviruses. While one could argue that the interaction with Tim-1 and Tim-4 is just one of many, soluble Tim-1-Fc protein exhibited a remarkable potency at inhibiting KSHV infection of different cells, and to a certain degree also inhibited KSHV infection of Tim-1-negative cells, whereas it did not affect entry of EBOV GP-pseudotyped lentiviral particles. This indicates that Tim-1 interacts with structures that perform critical functions during KSHV entry and not just promotes endocytosis of the particle.

## MATERIAL AND METHODS

### Cell culture, transfection, and lentiviral transduction

A549^70^ and Human embryonic kidney (HEK) 293T cells (ATCC) were kindly provided by the laboratory of Stefan Pöhlmann, German Primate Center—Leibniz Institute for Primate Research, Göttingen, Germany. SLK^71^ cells (RRID:CVCL_9569) were originally obtained from the (NIH AIDS Research and Reference Reagent program). Rhesus monkey fibroblasts (RF) were gently provided by the laboratory of Rüdiger Behr, German Primate Center—Leibniz Institute for Primate Research, Göttingen, Germany. If not stated otherwise, all cells were cultured in Dulbecco’s Modified Eagle Medium (DMEM), high glucose, GlutaMAX, 25mM HEPES (Thermo Fisher Scientific) supplemented with 10% fetal bovine serum (FBS) (Serana Europe GmbH), and 50μg/ml gentamycin (PAN Biotech) (D10). iSLK cells^72^ were maintained in D10 supplemented with 2.5μg/ml puromycin (InvivoGen) and 250μg/ml G418 (Carl Roth), for selection of iSLK carrying KSHV BACs 200µg/ml hygromycin was added.

For production of lentiviral particles, 10cm cell culture grade petri dishes of approximately 80% confluent 293T cells were transfected with 1.4μg pMD2.G (VSV-G envelope expressing plasmid, a gift from Didier Trono (Addgene plasmid #12259), 3.6μg psPAX2 (Gag-Pol expression construct, a gift from Didier Trono (Addgene plasmid #12260), and 5μg of lentiviral expression constructs (pLenti CMV Blast DEST (706–1), pLenti-CMV-Blast-TIM1-Strep, pLenti-CMV-Blast-TIM3-Strep, pLenti-CMV-Blast-TIM4-Strep, pLenti-CMV-Blast-NRP1-Strep, pLenti-CMV-Blast-TIM1-N114A-Strep, pLenti-CMV-Blast-TIM1-ND115AA-Strep, pLenti-CMV-Blast-TIM1-ND115DN-Strep, pLenti-CMV-Blast-TIM1-WF113AA-Strep, pLenti-CMV-Blast-TIM1-ΔO-gly-Strep, pLenti-CMV-Blast-TIM1-TransM1-Strep, pLenti-CMV-Blast-TIM1-TransM2-Strep) using PEI as described before^73^. The supernatant containing the pseudotyped lentiviral particles was harvested 2 to 3 days after transfection and filtered through 0.45μm CA membranes (Millipore). For transduction, lentivirus stocks were used at a 1:2 dilution. After 48-72h, the selection antibiotic blasticidin (Invivogen) was added to a final concentration of 10μg/ml. After 72h of initial selection, the blasticidin concentration was reduced to 5μg/ml and then to 1μg/ml to keep the cells in culture.

### Knockout screen for host factors

The human CRISPR (clustered regularly interspaced short palindromic repeats) “Brunello” lentiviral pooled library was purchased from Addgene. The library version in the lentiCRISPRv2 backbone was chosen, which is a ready-to-use lentiviral pooled library for CRISPR screening coding for Streptococcus pyogenes Cas9 and unique small guide (sg)RNAs that target more than 19 000 genes in the human genome. The library was amplified according to the manufacturer’s protocol. To produce the library in its lentivirus-vectored form, five T175 flasks of HEK293T cells were transfected with the library and the necessary packing plasmids. Briefly, for each flask 12.5 µg of library plasmid, 9 µg of psPAX2 (packaging plasmid, gag-pol expression construct), and 3.5 µg of VSV-G (envelope plasmid, VSV-G expression construct) were mixed in 1 mL of Opti-MEM and then 75 µL of Polyethylenimine (PEI) was added^73^. The mixture was incubated at room temperature for 30 minutes and then added to the cells. Between six and sixteen hours later, the medium was replaced with fresh D10. Three days post transfection, the cell culture supernatant was collected and filtered through a 0.45 a µm PVDF membrane filter.

5×10^6^ A549 cells were seeded in T175 flasks. Six hours post seeding cells were transduced with the lentiviruses “Brunello” library at a multiplicity of infection (MOI) of approx. 0.3 in 20ml D10 + 10µg/ml 1,5-Dimethyl-1,5-diazaundecamethylen-polymethobromid, Hexadimethrinbromid (Polybrene, Merk). Twenty-four hours post transduction, the cells were selected with 10µg/ml puromycin for three days. Transduced and selected A549 were then infected at MOI 0.1-0.2 with RRV-YFP virus (wt or dM); transduced and selected but not infected cells were used as negative control. Sixteen hours post infection the medium was replaced with fresh D10 and then replaced again to D20 (D10 with an additional 10% FBS) when most of the cells died. Due to the lytic replication of RRV, only few cells survived. The surviving cells were harvested and gDNA was extracted by PureLink Genomic DNA Mini Kit (Invitrogen). sgRNA cassettes were amplified by PCR and NGS adapters were added using a specific forward primer (TCGTCGGCAGCGTCAGATGTGTATAAGAGACAGTATCTTGTGGAAAGGACGAAA) and reverse primer (GTCTCGTGGGCTCGGAGATGTGTATAAGAGACAGATTCCCACTCCTTTCAAGACCTAG).

The sgRNA cassettes were amplified through polymerase chain reaction (PCR) using DreamTaq® polymerase (5U/µL, ThermoFisher, EP0702). To achieve optimal library coverage, we implemented a strategy that involved dividing gDNA products into several PCR reactions. For this purpose, the total gDNA volume of 115 µL was used, with specific concentrations for each sample: 220 ng/µL for RRV wt, 210 ng/µL for RRVdM, and 196.36 ng/µL for the control sample (A549 transduced with Brunello library but not infected). Subsequently, we performed a total of 23 PCR reactions, each utilizing 5 µL of the respective gDNA product as the template using the following conditions: 95°C for 3 min, followed by 18 cycles of 95°C for 30 sec, 61°C for 30 sec and 72°C for 20 sec. This was followed by a final elongation step at 72°C for 3 min.

The resulting PCR products were analysed by next-generation sequencing. Illumina HiSeq2500 High output run 100bp PE sequencing was performed by Macrogen Inc. Raw sequencing data were obtained for each sample and then analysed with the python software Model-based Analysis of Genome-wide CRISPR/Cas9 Knockout (MAGeCK) computational tool version 0.5.9.2 using default settings^74^. Genes from the lentiviral library screen, that were enriched in surviving cells were identified.

The top hundred enriched-sgRNA genes, from MAGeCK analysis were sorted according to their location and or function in the cell by the Functional Enrichment analysis tool (FunRich) software^29^ to arrive at a manageable set of membrane protein encoding genes for re-testing. 11 genes that were selected by FunRich and additionally by manual curation as well as NRP1 were then followed up upon. Four sgRNAs for each gene were designed using E-Crispr Design (e-crispr.org) under strict conditions ((only NGG PAM, only G as 5’ base, off-target tolerates many mismatches and ignores non-seed region, introns, purpose is knockout (only first 3 coding exons are allowed) and UTRs are excluded). The oligos were synthesised and ordered from IDT. The sgRNA oligos were cloned into lenti-CRISPRv2 vector and the inserted sequences were confirmed by Sanger sequencing (sequencing primer GACTATCATATGCTTACCGT). If cloning failed, the construct was not further analysed, the same was done if lentivirus production or transduction failed, so that not for all genes four sgRNAs were tested.

### Plasmids

TIM1 [Ref.: HQ412639.1], TIM3 [Ref.: NM_032782.5], TIM4 [Ref.: NM_138379.3] (kindly provided by Stefan Pöhlmann’s laboratory, infection biology, DPZ, Germany) and NRP1 [Ref.: NM_003873.7] (kindly provided by the laboratory of Mikael Simons, DZNE, Munich), followed by the Strep-Tag (OneStrep Tag: SAWSHPQFEKGGGSGGGSGGSAWSHPQFEK), were cloned into CMV Blast DEST (706–1) (a gift from Eric Campeau & Paul Kaufman, Addgene plasmid #17451) expression vector resulting in plasmid AX581. TIM1 mutants (N114A, ND115AA, ND115DN, WF113AA, ΔO-glyc) and TIM1 with truncated cytoplasmatic domain (TIM1_334^STOP^ and TIM1_311^STOP^, termed TIM1_ΔIC) constructs were generated based on AX581 using ‘Round the Horn’ Site-directed mutagenesis^75^. The soluble ectodomain (amino acids 1-290) of Tim-1 was C-terminally fused to IgG1 Fc with a C-terminal tandem Strep-Tag and 6x His tag in the pAB61 vector^6^ by Gibson assembly. All constructs were verified by Sanger sequencing.

### BAC mutagenesis

Mutant viruses, used in the present study, were generated using a two-step, markerless λ-red-mediated BAC recombination strategy as described by Tischer et al. [109]. Briefly, recombination cassettes were generated from the pEPKanS template with Phusion High Fidelity DNA polymerase (Thermo Fisher Scientific) by polymerase chain reaction (PCR) using long oligonucleotides (Ultramers; purchased from Integrated DNA Technologies, IDT). Recombination cassettes were transformed into BAC16-carrying Escherichia coli strain GS1783 or RRV-YFP-carrying GS1783 respectively, followed by kanamycin selection, and subsequent second recombination under 1% L(+)arabinose (Sigma-Aldrich)-induced I-SceI expression. Colonies were verified by PCR of the mutated region followed by sequence analysis (Macrogen), pulsed-field gel electrophoresis and restriction fragment length polymorphism. For this purpose, bacmid DNA was isolated by standard alkaline lysis from 5ml liquid cultures. Subsequently, the integrity of bacmid DNA was analysed by digestion with restriction enzyme XhoI and separation in 0.8% PFGE agarose (Bio-Rad) gels and 0.5×TBE buffer by pulsed-field gel electrophoresis at 6 V/cm, 120-degree field angle, switch time linearly ramped from 1s to 5s over 16 h (CHEF DR III, Bio-Rad).

### Virus production and infection experiments

Infection assay were performed plating HEK 293T cells at 40000 cells/well and A549 or SLK at 25000 cells/well in 96-well plates. 24 hours post plating, the cells were infected with the indicated amounts of virus. One day post infection, cells were harvested by brief trypsinisation, followed by addition of 5% FBS in PBS to inhibit trypsin activity. Then, the cell suspension was transferred to an equal volume of PBS supplemented with 4% methanol-free formaldehyde (Carl Roth) for fixation. A minimum of 5000 cells/well were analysed for GFP and/or YFP expression by Sony ID7000 spectral cell analyser (Sony™). For blocking experiments, soluble Fc decoy receptors were incubated for thirty minutes at room temperature with the respective viruses at 30μg/ml, unless stated otherwise. After the incubation period, selected cells were infected in 96-well plates as described above.

For blocking experiments with anti-Tim-1 antibody, the cells were preincubated with antibody or isotype control for 30 minutes. Then virus was added in a small volume (final concentration of antibody after addition of virus). IgG1 isotype antibody was purchased from InVivoMAb (clone MOPC 21, Cat# BE0083), ARD5 anti-Tim-1 antibody was cloned according to the sequence kindly provided by Wendy Maury with mouse IgG1 heavy chain, produced by Genscript and provided in PBS^41^.

### Attachment assay

The attachment assay was performed as described previously^25^. Briefly, cells were incubated with KSHV and spin-inoculated by centrifugation at 4°C and 4122g for 30min. After three washes with ice-cold PBS, genomic DNA was isolated using the ISOLATE II Genomic DNA Kit (Bioline) according to the manufacturer’s instructions. The amount of viral DNA was determined by qPCR from equal amounts of extracted DNA.

### Recombinant protein production

Recombinant, soluble FcStrep, Tim1-Fc and mutant Tim-1-Fc proteins were purified by Strep-Tactin chromatography from HEK 293T cell culture supernatant. HEK 293T cells were transfected with the respective expression plasmids. The protein-containing cell culture supernatant was filtered through 0.22μm PES membranes (Millipore) and passed over 0.5ml of a Strep-Tactin Superflow (IBA Lifesciences) matrix in a gravity flow Omniprep column (BioRad). Bound protein was washed with approximately 50ml phosphate buffered saline pH 7.4 (PBS) and eluted in 1ml fractions with 3mM desthiobiotin (Sigma-Aldrich) in PBS, alternatively also with 50mM Biotin in 150mM NaCl 50mM Tris pH 7.5. Protein-containing fractions were pooled, and buffer exchange to PBS using VivaSpin columns (Sartorius) was carried out. The absorbance at 280 nm was used to estimate the protein concentration. For storage, aliquots were frozen and kept at −80°C.

### Immunofluorescence

Immunofluorescence was performed as described previously^52^. SLK cells were seeded at approx. 75 000 cells per well on 12-mm coverslips (YX03.1; Carl Roth) in 24-well plates. On day two, the plate was cooled down on ice, followed by inoculation with 1000µl cold virus suspension per cell and 30min centrifugation at 4122g and 4°C, followed by another 10min at 4°C. Then, the cells were washed three times in ice-cold PBS, followed either by fixation or medium exchange to D10 and incubation for 2h at 37°C in a cell culture incubator, followed by one wash in PBS and fixation. The cells were fixed in 4% methanol-free formaldehyde in PBS for 10 min. After fixation, the cells were permeabilized in IF buffer (PBS supplemented with 5% FBS and 0.05% Saponin) for 1h at room temperature, incubated with 40µl primary antibody (anti-Tim-1, 1:500) for 2h in IF buffer in a humid chamber, followed by three washes with IF buffer, followed by incubation with secondary antibody for 1h at room temperature, otherwise identical as for the primary antibody, followed by one wash in IF buffer, followed by one wash in PBS supplemented with Hoechst dye, followed by one wash in PBS. The coverslips were then mounted in antifade mounting medium (Abcam).

Z-stacks were imaged using ZEN software (Zeiss) and a laser scanning microscope (Zeiss SLM 800), frame size 2048px x 2048px, scan speed 5, pinhole 58µm, interval 0.31µm, one separate scan for the AlexaFluor647 channel, one for the EGFP channel, and one for the Hoechst channel. Two similar planes that captured the middle of the cell layer from each stack were used for analysis, three independent sets of experiments were analysed using the FIJI (https://fiji.sc/) edition of ImageJ. Colocalized particles were identified using the plugin “Colocalization Finder” by Pilippe Carl. Limits were set so that no colocalisation was observed in controls without virus or Tim-1 primary antibody. The region of interest containing the colocalized signals was then copied to the original image, and viral particles within this ROI were the quantified using the “find maxima” function of ImageJ in the green channel image, the same was also done for the whole image. The ratio of colocalized particles over all particles was then calculated.

## ACKNOWLEDGEMENTS

We thank Stefan Pöhlmann for support, Wendy Maury for the ARD5 sequence Christian Münz for helpful discussions, Rüdiger Behr for rhesus monkey fibroblasts, Klaus Korn for human foreskin fibroblasts.

This work was funded by grants to ASH from the Deutsche Forschungsgemeinschaft (grants HA 6013/4-1, DFG HA 6013/10-1), by an Exploration Grant from the Boehringer-Ingelheim Foundation to A.S.H., by a grant from the Wilhelm-Sander Foundation (https://www.wilhelm-sander-stiftung.de; project no. 2019.027.1) to ASH. X.Y. was funded by China Petroleum Central Hospital. S.L. was funded by a scholarship of the China Scholarship Council (CSC), file number 202106300006.

## Notes

### Competing Interest Statement

Alexander S. Hahn is also an employee of GSK. GSK had no influence on the study.

### Summary of Updates

Inhibition experiments using a monoclonal antibody to Tim-1 have been added. Minor corrections were made.

## Reference

1. Cesarman, E. et al. Kaposi sarcoma. Nat Rev Dis Primers 5, 1–21 (2019).

2. Bray, F. et al. Global cancer statistics 2018: GLOBOCAN estimates of incidence and mortality worldwide for 36 cancers in 185 countries. CA Cancer J Clin (2018) doi:10.3322/caac.21492.

3. Chen, Q., et al. Kaposi’s sarcoma herpesvirus is associated with osteosarcoma in Xinjiang populations. PNAS 118, (2021).

4. Wen, K. W., Wang, L., Menke, J. R. & Damania, B. Cancers associated with human gammaherpesviruses. The FEBS Journal 289, 7631–7669 (2022).

5. Goncalves, P. H., Ziegelbauer, J., Uldrick, T. S. & Yarchoan, R. Kaposi sarcoma herpesvirus-associated cancers and related diseases. Current Opinion in HIV and AIDS 12, 47–56 (2017).

6. Birkmann, A. et al. Cell Surface Heparan Sulfate Is a Receptor for Human Herpesvirus[8 and Interacts with Envelope Glycoprotein K8.1. Journal of Virology 75, 11583–11593 (2001).

7. Hahn, A. et al. Kaposi’s sarcoma-associated herpesvirus gH/gL: glycoprotein export and interaction with cellular receptors. J Virol 83, 396–407 (2009).

8. Garrigues, H. J., DeMaster, L. K., Rubinchikova, Y. E. & Rose, T. M. KSHV attachment and entry are dependent on αVβ3 integrin localized to specific cell surface microdomains and do not correlate with the presence of heparan sulfate. Virology 0, 118–133 (2014).

9. Akula, S. M., Pramod, N. P., Wang, F.-Z. & Chandran, B. Integrin α3β1 (CD 49c/29) Is a Cellular Receptor for Kaposi’s Sarcoma-Associated Herpesvirus (KSHV/HHV-8) Entry into the Target Cells. Cell 108, 407–419 (2002).

10. Garrigues, H. J., Rubinchikova, Y. E., DiPersio, C. M. & Rose, T. M. Integrin {alpha}V 3 Binds to the RGD Motif of Glycoprotein B of Kaposi’s Sarcoma-Associated Herpesvirus and Functions as an RGD-Dependent Entry Receptor. J. Virol. 82, 1570–1580 (2008).

11. Hahn, A. S. et al. The ephrin receptor tyrosine kinase A2 is a cellular receptor for Kaposi’s sarcoma–associated herpesvirus. Nat Med 18, 961–966 (2012).

12. Großkopf, A. K. et al. A conserved Eph family receptor-binding motif on the gH/gL complex of Kaposi’s sarcoma-associated herpesvirus and rhesus monkey rhadinovirus. PLoS Pathog. 14, e1006912 (2018).

13. Hahn, A. S. & Desrosiers, R. C. Binding of the Kaposi’s sarcoma-associated herpesvirus to the ephrin binding surface of the EphA2 receptor and its inhibition by a small molecule. J Virol 88, 8724–8734 (2014).

14. Hahn, A. S. & Desrosiers, R. C. Rhesus Monkey Rhadinovirus Uses Eph Family Receptors for Entry into B Cells and Endothelial Cells but Not Fibroblasts. PLOS Pathogens 9, e1003360 (2013).

15. van der Meulen, E., Anderton, M., Blumenthal, M. J. & Schäfer, G. Cellular Receptors Involved in KSHV Infection. Viruses 13, 118 (2021).

16. Inoue, N., Winter, J., Lal, R. B., Offermann, M. K. & Koyano, S. Characterization of Entry Mechanisms of Human Herpesvirus 8 by Using an Rta-Dependent Reporter Cell Line. J Virol 77, 8147–8152 (2003).

17. Akula, S. M. et al. Kaposi’s Sarcoma-Associated Herpesvirus (Human Herpesvirus 8) Infection of Human Fibroblast Cells Occurs through Endocytosis. Journal of Virology 77, 7978–7990 (2003).

18. Raghu, H., Sharma-Walia, N., Veettil, M. V., Sadagopan, S. & Chandran, B. Kaposi’s sarcoma-associated herpesvirus utilizes an actin polymerization-dependent macropinocytic pathway to enter human dermal microvascular endothelial and human umbilical vein endothelial cells. J. Virol. 83, 4895–4911 (2009).

19. TerBush, A. A., Hafkamp, F., Lee, H. J. & Coscoy, L. A Kaposi’s Sarcoma-Associated Herpesvirus Infection Mechanism is Independent of Integrins α3β1, αVβ3, and αVβ5. J. Virol. (2018) doi:10.1128/JVI.00803-18.

20. Großkopf, A. K. et al. EphA7 Functions as Receptor on BJAB Cells for Cell-to-Cell Transmission of the Kaposi’s Sarcoma-Associated Herpesvirus and for Cell-Free Infection by the Related Rhesus Monkey Rhadinovirus. J Virol 93, e00064–19 (2019).

21. Fricke, T., Großkopf, A. K., Ensser, A., Backovic, M. & Hahn, A. S. Antibodies Targeting KSHV gH/gL Reveal Distinct Neutralization Mechanisms. Viruses 14, 541 (2022).

22. Großkopf, A. K. et al. Plxdc family members are novel receptors for the rhesus monkey rhadinovirus (RRV). PLoS Pathog 17, e1008979 (2021).

23. Dollery, S. J., Santiago-Crespo, R. J., Chatterjee, D. & Berger, E. A. Glycoprotein K8.1A of Kaposi’s sarcoma-associated herpesvirus is a critical B cell tropism determinant, independent of its heparan sulfate binding activity. J. Virol. (2018) doi:10.1128/JVI.01876-18.

24. Hahn, A. S. et al. A Recombinant Rhesus Monkey Rhadinovirus Deleted of Glycoprotein L Establishes Persistent Infection of Rhesus Macaques and Elicits Conventional T Cell Responses. J Virol 94, e01093–19 (2020).

25. Liu, S. et al. Kaposi’s sarcoma-associated herpesvirus glycoprotein K8.1 is critical for infection in a cell-specific manner and functions at the attachment step on keratinocytes. 2023.03.19.533316 Preprint at 10.1101/2023.03.19.533316 (2023).

26. Doench, J. G. et al. Optimized sgRNA design to maximize activity and minimize off-target effects of CRISPR-Cas9. Nat. Biotechnol. 34, 184–191 (2016).

27. The Human Protein Atlas. https://www.proteinatlas.org/.

28. Uhlén, M. et al. Proteomics. Tissue-based map of the human proteome. Science 347, 1260419 (2015).

29. Pathan, M. et al. FunRich: An open access standalone functional enrichment and interaction network analysis tool. Proteomics 15, 2597–2601 (2015).

30. Kobayashi, N. et al. TIM-1 and TIM-4 Glycoproteins Bind Phosphatidylserine and Mediate Uptake of Apoptotic Cells. Immunity 27, 927–940 (2007).

31. Freeman, G. J., Casasnovas, J. M., Umetsu, D. T. & DeKruyff, R. H. TIM genes: a family of cell surface phosphatidylserine receptors that regulate innate and adaptive immunity. Immunol Rev 235, 172–189 (2010).

32. Kaplan, G. et al. Identification of a surface glycoprotein on African green monkey kidney cells as a receptor for hepatitis A virus. EMBO J 15, 4282–4296 (1996).

33. Brouillette, R. B. et al. TIM-1 Mediates Dystroglycan-Independent Entry of Lassa Virus. J Virol 92, e00093–18 (2018).

34. The Phosphatidylserine Receptor TIM-1 Enhances Authentic Chikungunya Virus Cell Entry - PubMed. https://pubmed.ncbi.nlm.nih.gov/34359995/.

35. Niu, J. et al. TIM-1 Promotes Japanese Encephalitis Virus Entry and Infection. Viruses 10, 630 (2018).

36. Moller-Tank, S., Albritton, L. M., Rennert, P. D. & Maury, W. Characterizing functional domains for TIM-mediated enveloped virus entry. J Virol 88, 6702–6713 (2014).

37. Zhang, X. et al. T-Cell Immunoglobulin and Mucin Domain 1 (TIM-1) Is a Functional Entry Factor for Tick-Borne Encephalitis Virus. mBio 13, e0286021 (2022).

38. Meertens, L. et al. The TIM and TAM families of phosphatidylserine receptors mediate dengue virus entry. Cell Host Microbe 12, 544–557 (2012).

39. Morizono, K. & Chen, I. S. Y. Role of phosphatidylserine receptors in enveloped virus infection. J Virol 88, 4275–4290 (2014).

40. Wang, J., Qiao, L., Hou, Z. & Luo, G. TIM-1 Promotes Hepatitis C Virus Cell Attachment and Infection. J Virol 91, e01583–16 (2017).

41. Kondratowicz, A. S. et al. T-cell immunoglobulin and mucin domain 1 (TIM-1) is a receptor for Zaire Ebolavirus and Lake Victoria Marburgvirus. Proc Natl Acad Sci U S A 108, 8426–8431 (2011).

42. Wang, H.-B. et al. Neuropilin 1 is an entry factor that promotes EBV infection of nasopharyngeal epithelial cells. Nat Commun 6, 6240 (2015).

43. Lu, Z.-Z. et al. Neuropilin 1 is an entry receptor for KSHV infection of mesenchymal stem cell through TGFBR1/2-mediated macropinocytosis. Sci Adv 9, eadg1778 (2023).

44. Moller-Tank, S., Kondratowicz, A. S., Davey, R. A., Rennert, P. D. & Maury, W. Role of the Phosphatidylserine Receptor TIM-1 in Enveloped-Virus Entry. J Virol 87, 8327– 8341 (2013).

45. Santiago, C. et al. Structures of T Cell immunoglobulin mucin receptors 1 and 2 reveal mechanisms for regulation of immune responses by the TIM receptor family. Immunity 26, 299–310 (2007).

46. Moon, B. et al. EphA2 Interacts with Tim-4 through Association between Its FN3 Domain and the IgV Domain of Tim-4. Cells 10, 1290 (2021).

47. Chen, J., Zhang, X., Schaller, S., Jardetzky, T. S. & Longnecker, R. Ephrin Receptor A4 is a New Kaposi’s Sarcoma-Associated Herpesvirus Virus Entry Receptor. mBio 10, e02892–18 (2019).

48. Su, C. et al. Molecular basis of EphA2 recognition by gHgL from gammaherpesviruses. Nat Commun 11, 5964 (2020).

49. Kaleeba, J. A. R. & Berger, E. A. Kaposi’s Sarcoma-Associated Herpesvirus Fusion-Entry Receptor: Cystine Transporter xCT. Science 311, 1921–1924 (2006).

50. Veettil, M. V. et al. Kaposi’s Sarcoma-Associated Herpesvirus Forms a Multimolecular Complex of Integrins (αVβ5, αVβ3, and α3β1) and CD98-xCT during Infection of Human Dermal Microvascular Endothelial Cells, and CD98-xCT Is Essential for the Postentry Stage of Infection. Journal of Virology 82, 12126–12144 (2008).

51. Dutta, D. et al. EphrinA2 Regulates Clathrin Mediated KSHV Endocytosis in Fibroblast Cells by Coordinating Integrin-Associated Signaling and c-Cbl Directed Polyubiquitination. PLoS Pathog 9, e1003510 (2013).

52. Hörnich, B. F., Großkopf, A. K., Dcosta, C. J., Schlagowski, S. & Hahn, A. S. Interferon-Induced Transmembrane Proteins Inhibit Infection by the Kaposi’s Sarcoma-Associated Herpesvirus and the Related Rhesus Monkey Rhadinovirus in a Cell-Specific Manner. mBio 12, e0211321 (2021).

53. Saeed, M. F., Kolokoltsov, A. A., Albrecht, T. & Davey, R. A. Cellular entry of ebola virus involves uptake by a macropinocytosis-like mechanism and subsequent trafficking through early and late endosomes. PLoS Pathog 6, e1001110 (2010).

54. Ding, Q. et al. Regulatory B cells are identified by expression of TIM-1 and can be induced through TIM-1 ligation to promote tolerance in mice. J Clin Invest 121, 3645–3656 (2011).

55. Ma, J. et al. TIM-1 signaling in B cells regulates antibody production. Biochem Biophys Res Commun 406, 223–228 (2011).

56. Zhang, Y. et al. CD19+ Tim-1+ B cells are decreased and negatively correlated with disease severity in Myasthenia Gravis patients. Immunol Res 64, 1216–1224 (2016).

57. Wong, K. et al. Phosphatidylserine receptor Tim-4 is essential for the maintenance of the homeostatic state of resident peritoneal macrophages. Proc Natl Acad Sci U S A 107, 8712–8717 (2010).

58. Bruce, A. G. et al. Macaque Homologs of EBV and KSHV Show Uniquely Different Associations with Simian AIDS-related Lymphomas. PLoS Pathog. 8, e1002962 (2012).

59. McHugh, D. et al. Persistent KSHV Infection Increases EBV-Associated Tumor Formation In Vivo via Enhanced EBV Lytic Gene Expression. Cell Host Microbe 22, 61–73.e7 (2017).

60. Shimada, K., Hayakawa, F. & Kiyoi, H. Biology and management of primary effusion lymphoma. Blood 132, 1879–1888 (2018).

61. Myoung, J. & Ganem, D. Infection of primary human tonsillar lymphoid cells by KSHV reveals frequent but abortive infection of T cells. Virology 413, 1–11 (2011).

62. Subbarayal, P. et al. EphrinA2 Receptor (EphA2) Is an Invasion and Intracellular Signaling Receptor for Chlamydia trachomatis. PLoS Pathog 11, e1004846 (2015).

63. Lupberger, J. et al. EGFR and EphA2 are host factors for hepatitis C virus entry and possible targets for antiviral therapy. Nat Med 17, 589–595 (2011).

64. Dong, X.-D. et al. EphA2 is a functional entry receptor for HCMV infection of glioblastoma cells. PLOS Pathogens 19, e1011304 (2023).

65. Zhang, H. et al. Ephrin receptor A2 is an epithelial cell receptor for Epstein-Barr virus entry. Nat Microbiol (2018) doi:10.1038/s41564-017-0080-8.

66. Aaron, P. A., Jamklang, M., Uhrig, J. P. & Gelli, A. The blood-brain barrier internalises Cryptococcus neoformans via the EphA2-tyrosine kinase receptor. Cell. Microbiol. 20, (2018).

67. Bohan, D. et al. Phosphatidylserine receptors enhance SARS-CoV-2 infection. PLoS Pathog 17, e1009743 (2021).

68. Sui, L. et al. Human membrane protein Tim-3 facilitates hepatitis A virus entry into target cells. Int J Mol Med 17, 1093–1099 (2006).

69. Ashida, M. & Hamada, C. Molecular cloning of the hepatitis A virus receptor from a simian cell line. J Gen Virol 78 **( Pt** **7**), 1565–1569 (1997).

70. Giard, D. J. et al. In vitro cultivation of human tumors: establishment of cell lines derived from a series of solid tumors. J Natl Cancer Inst 51, 1417–1423 (1973).

71. Stürzl, M., Gaus, D., Dirks, W. G., Ganem, D. & Jochmann, R. Kaposi’s sarcoma-derived cell line SLK is not of endothelial origin, but is a contaminant from a known renal carcinoma cell line. Int. J. Cancer 132, 1954–1958 (2013).

72. Myoung, J. & Ganem, D. Generation of a doxycycline-inducible KSHV producer cell line of endothelial origin: Maintenance of tight latency with efficient reactivation upon induction. Journal of Virological Methods 174, 12–21 (2011).

73. Longo, P. A., Kavran, J. M., Kim, M.-S. & Leahy, D. J. Transient Mammalian Cell Transfection with Polyethylenimine (PEI). Methods Enzymol 529, 227–240 (2013).

74. Li, W. et al. MAGeCK enables robust identification of essential genes from genome-scale CRISPR/Cas9 knockout screens. Genome Biology 15, 554 (2014).

75. Hörnich, B. F. et al. SARS-CoV-2 and SARS-CoV Spike-Mediated Cell-Cell Fusion Differ in Their Requirements for Receptor Expression and Proteolytic Activation. J Virol 95, e00002–21 (2021).

